# Testing fundamental hypotheses of colonization success in the ferns

**DOI:** 10.64898/2026.04.16.719039

**Authors:** Jessie A. Pelosi, Agustina Yañez, Leah N. Veldhuisen, Anthony J. Dant, Poppy C. Northing, Rebecca G.W. Bland, Weston L. Testo, Katrina M. Dlugosch

## Abstract

**Background and Aims:** Non-native species are now ubiquitous members of regional floras. The factors that lead to establishment and dominance of non-native species are continuously debated. Fundamental hypotheses about drivers of invasion success include the role of phylogeny, polyploidy, genome size, and rapid niche evolution. These hypotheses have been tested in the seed plants, but ferns, the second largest group of vascular plants, have rarely been considered in these analyses, despite making up a non-trivial portion of non-native floras.

**Methods:** We compiled a dataset of global non-native ferns and categorized them along the invasion spectrum using descriptions from the literature and natural history collections. Using this dataset, we assessed I) the taxonomic diversity and phylogenetic clustering of non-native ferns, II) the geographic distribution of fern introductions, testing for shifts in climatic niches, and III) test for the association of ‘invader’ traits across the invasion continuum, including smaller genome sizes and higher ploidal levels.

**Key Results:** We generated a dataset that includes 83 taxa; of these, we classified 18 as casual, 35 as naturalized (but not invasive), and 30 as invasive. Using this dataset, we found I) weak or no phylogenetic clustering of non-native ferns, II) some regions are overrepresented as sources and recipients of introductions, III) climatic niches are often conserved between native and introduced ranges, but can differ between introductions, IV) naturalized ferns have smaller genomes, and V) invaders have higher ploidal levels.

**Conclusions:** We integrated regional floras, occurrence and climate data, phylogeny, and cytology to test fundamental hypotheses regarding the colonization success of ferns. This study provides insights into the ecological, genomic, and phylogenetic features associated with the colonization of new habitats by non-native ferns, a largely overlooked portion of non-native plant taxa.

## INTRODUCTION

Non-native taxa are now ubiquitous members of the world’s flora (Vellend *et al*. 2013), and those that form established, self-sustaining populations can have substantial economic and ecological impacts (Pyšek *et al*. 2020). Non-native species that become dominant members of their communities can alter native species’ abundance and risk of extinction (Wilcove *et al*. 1998; Mooney and Cleland 2001), community richness and composition (Vilà *et al*. 2011), and ecosystem biogeochemical cycles and productivity (Gaertner *et al*. 2014; Capinha *et al*. 2015). As a result, the factors that lead to establishment and dominance of non-native species have been under intense investigation since species invasions became a broadly recognized phenomenon (Elton 1958; Baker and Stebbins 1965; Richardson 2011).

The factors that facilitate invasions have been studied from the perspective of species (Pyšek and Richardson 2007; Pyšek *et al*. 2014; van Kleunen *et al*. 2016), invaded regions or environments (Lonsdale 1999; Richardson and Pyšek 2012; Deslippe and Veenendaal 2025), introduction agents (McCulloch-Jones *et al*. 2021) or particularities of the introduction event (Catford *et al*. 2009; Simberloff 2009; Maurel *et al*. 2016; Pyšek *et al*. 2020; Gioria *et al*. 2023). Despite the complexity of the factors involved in this process, invasion biologists generally agree that there exists a series of stages or phases in which a non-native species establishes and spreads in a new area: the introduction-naturalization-invasion continuum, frequently termed simply the invasion continuum (Richardson *et al*. 2000; Colautti and MacIsaac 2004; Blackburn *et al*. 2011). To progress through the stages of the invasion continuum after introduction, a species must overcome environmental and reproductive barriers, and a variety of hypotheses (Enders *et al*. 2020) have been proposed to explain how successful invaders do so (e.g. Williamson and Fitter 1996; Richardson *et al*. 2000; Pyšek *et al*. 2004; Falk-Petersen *et al*. 2006).

To date, most tests about the invasion biology of plants have been focused on the seed plants (e.g., van Kleunen *et al*. 2016). Ferns are the second largest group of vascular plants (>10,000 species, PPG I 2016), and yet are largely overlooked in their contributions to non-native floras. While ferns are commonly associated with a humid, low-light, terrestrial environment, they occupy a wide variety of ecological niches across the globe, show very different forms, growth habits, traits, and reproductive strategies, and are responsible for some of the most damaging invasions globally (Pemberton and Ferriter 1998; Robinson *et al*. 2010). In addition, since ferns diverged from the seed plants over 400 million years ago, they provide an excellent opportunity to test fundamental hypotheses in invasion biology in a distinct evolutionary context (Testo and Sundue 2016; Morris *et al*. 2018).

Most non-native ferns have been introduced through horticulture and cultivation (Robinson *et al*. 2010; McCulloch-Jones *et al*. 2023), eventually escaping and becoming self-sustaining populations. Currently, very few authors have considered ferns in their analyses of non-native taxa (Robinson *et al*. 2010; Akomolafe and Rahmad 2018; Jones *et al*. 2019). The most recent assessment of non-native ferns by (Jones *et al*. 2019) tested whether functional traits and habitat preferences could explain invasiveness. Their model predicted that high reproductive plasticity and ability to tolerate various disturbance and light conditions were strongly positively correlated with invasion success in terrestrial ferns. As Jones *et al*. (2019) note, other intrinsic (e.g., traits) and extrinsic (e.g., ecological and evolutionary processes) features also need to be considered when determining what drives invasion success in ferns.

In this article, we assemble a new global catalog of non-native ferns and identify their position along the invasion continuum by drawing on documented observations in regional floras and the literature. We use this new catalog to describe non-native fern diversity and to test major hypotheses for the success of non-native species in ferns. Given the current knowledge gap regarding ferns as non-native species, our study focuses primarily on the intrinsic properties that could facilitate their invasion. We ask whether non-native ferns that successfully establish possess distinctive traits, and whether there are key biological or ecological characteristics that can predict why some species progress beyond naturalization to become invasive. Specifically, in this study we: 1) describe the taxonomic diversity of non-native ferns and test for phylogenetic clustering of fern introductions and invasions (Pyšek 1998), 2) describe the geographic distribution of fern introductions, test for over- and underrepresentation of global regions as sources and destinations of introductions (van Kleunen *et al*. 2015), and test for shifts in the climatic niches of an example subset of invasive ferns (Petitpierre *et al*. 2012), and 3) test for the association of “invader” traits with the invasion continuum in ferns, including smaller genome size and higher ploidy level (Pandit *et al*. 2014). Each of these tests addresses a standing hypothesis in the invasion biology literature (see Discussion for further background). Through these analyses, we aim to provide insights into the ecological, genomic, and phylogenetic features associated with the colonization of new habitats by non-native ferns.

## METHODS

### Dataset Compilation

Our dataset of non-native ferns was developed by first downloading the World Ferns database (Hassler 2021, last accessed 11 May 2021) and sorting for introduced taxa. This dataset was supplemented with data from the Global Naturalized Alien Flora (GloNAF, van Kleunen *et al*. 2019) and Appendix I from Jones *et al*. (2019). Synonyms for taxa not included in the World Ferns database were determined using the World Flora Online (WFO 2021). We filtered the combined list of non-native taxa by first verifying the presence of non-native taxa with regional and local floras and/or peer-reviewed publications (Table S1). The geographic range of each taxon was also determined using the World Ferns database (Hassler, 2021), peer-reviewed publications, and occurrence data from Suissa *et al*. (2021; ∼900k expert-verified occurrence points).

Non-native taxa were classified according to their status on the invasion continuum Richardson *et al*. (2000), Colautti and MacIsaac (2004), and Blackburn *et al*. (2011), using a modified scheme from Yañez *et al*. (2023), as being “non-native”, “casual/adventive” (from now on “casual”), or “naturalized, not invasive” (from now on “naturalized”), or “naturalized, invasive” (from now on “invasive”). Species were assigned to this last category (“invasive”) if they produce viable offspring in large numbers and disperse over significant distances from the parent plants. This stage indicates a high potential for broad spatial expansion (Richardson et al. 2000; Pyšek et al. 2004; Richardson & Pyšek 2012) and ecological dominance within recipient communities (Table 1). We recorded quotes from the primary literature, floras, or herbarium labels to inform and justify our classifications (Table S1). We were primarily focused on identifying information about the abundance and dominance in the community to differentiate between the “naturalized” and “invasive” categories. Finally, if a species exhibited different stages of invasion across countries or regions, it was assigned to the most advanced stage observed within its introduced range. This approach reflects the species’ potential to become naturalized or invasive elsewhere.

**Table 1.** Terminology used in this study related to the introduction and status of non-native species. Table modified from Yañez *et al*. (2023).

| Term | Definition | References |
| --- | --- | --- |
| Native | Plant taxa that have originated in a given area without human involvement or that have arrived there without intentional or unintentional interventions of humans from an area in which they are native. | Pyšek *et al.* (2004); Falk-Petersen *et al.* (2006); Colauti & MacIssac (2004) |
| Non-native | Plant taxa in a given area, outside its natural past of present range, whose presence there is due to intentional or unintentional human involved, or which have arrived there without the help of people from an area in which they are non-native | Richardson *et al.* (2000); Pyšek *et al.* (2004); Falk-Petersen *et al.* (2006); Colauti & MacIssac (2004) |
| Cultivated | Plant taxa growing under human control, whose sustainability depends on humans | Pyšek *et al.* (2004) |
| Causal or Adventive (Stage II *sensu* Colauti & MacIssac, 2004) | Non-native plants that may flourish and even reproduce occasionally outside cultivation in an area, but that eventually die out because they do not form self-sustaining populations, and rely on repeated introductions for their persistence. | Richardson *et al.* (2000); Pyšek *et al.* (2004); Falk-Petersen *et al.* (2006); Colauti & MacIssac (2004) |
| Naturalized or non-invasive naturalized (Stage III *sensu* Colauti & MacIssac, 2004) | Non-native plants that reproduce consistently and have achieved a self-sustaining population over many generations without direct intervention by humans (or despite human intervention). Do not necessarily invade natural, seminatural, or human-made ecosystems and do not spread substantially. | Richardson *et al.* (2000); Pyšek *et al.* (2004); Falk-Petersen *et al.* (2006); Colauti & MacIssac (2004) |
| Invasive or invasive naturalized (Stage V *sensu* Colauti & MacIssac, 2004) | Non-native plants that reproduce consistently and have obtained a self-sustaining population over many generations without direct intervention by humans. These species become dominant members in the community and have the potential to spread over long distances. | Richardson & Pyšek (2012); Colauti & MacIssac (2004) |

### Phylogenetic Distribution

To determine the phylogenetic distribution of invasive ferns, we used the phylogeny from Fern Tree of Life (FTOL) v1.7.0 (Nitta *et al*. 2022). One species was present in FTOL under a synonym and two species were not present (Table S1). For the latter, we substituted the closest relative, usually a sister species. The pattern of phylogenetic clustering of binary traits can be modeled using the *D* statistic (Fritz and Purvis 2010), where values of *D* near 0 indicate trait conservatism consistent with Brownian motion, while values near 1 indicate random evolution. Values either below 0 or above 1 represent highly conserved or overdispersed traits, respectively (Orme *et al*. 2023). We calculated *D* with three different cutoffs for invasiveness: the least restrictive, where we scored casual, naturalized and invasive as a 1 and native as a 0, the moderately restrictive, where we scored naturalized and invasive as a 1 and casual and native as a 0, and the most restrictive, where we only scored invasive as a 1 and all others as a 0. We calculated the *D* statistic in the R package ‘caper’ v1.0.03 (Orme *et al*. 2023) with the function ‘phylo.d(),’ which gives one *P*-value based on Brownian motion and one based on random association.

We also calculated phylogenetic signal for invasiveness as a continuous trait with Blomberg’s K (Blomberg *et al*. 2003) and Pagel’s λ (Pagel 1999). We assigned numbers to each category (0: native, 1: casual, 2: naturalized, 3: invasive). We calculated Blomberg’s K with the “phylosignal()” function in the picante v1.8.2 R package (Kembel et al., 2010) with 5000 randomizations, and Pagel’s λ with the function “phylosig()” in the phytools v2.4-4 R package (Revell 2012) with 5000 iterations and model set to “lambda.” While phylogeny size should not impact results (Fritz & Purvis, 2010), we also pruned FTOL to only include the 82 non-native ferns, and recalculated Blomberg’s K (Blomberg *et al*. 2003) and Pagel’s λ (Pagel 1999). To identify which taxa/clades contribute to phylogenetic signal/clustering, we computed the Local Indicator of Phylogenetic Association with the function ‘lipaMoran’ in the phylosignal v1.3.1 (Keck *et al*. 2016) R package with 999 permutations using the entire FTOL and invasiveness as a continuous trait. We noted taxa that were significantly more similar to their closest relative than expected by chance (Ii > 0 and P < 0.05).

### Geographic Distribution

We designated a total of nine global regions, slightly modified from the regions defined by Moran (2008), and categorized the native and introduced range of each taxon by these regions. We visualized species movement between and within geographic regions with chord diagrams using the R package circlize v0.4.11 (Gu *et al*. 2014). We then used χ^2^ tests to determine if particular regions were overrepresented as sources or recipients of non-native species than expected. We estimated the number of taxa within each region using only the occurrence records from Suissa *et al*. (2021). Where occurrence lists indicate multiple native regions, we included all as potential source populations for introductions. Over-represented regions were identified as those with χ^2^ residuals > 2, and under-represented regions were identified as those with χ^2^ residuals < -2.

### Climatic Niche Shifts

To address how native and introduced range climatic niches compare and whether there are shifts between ranges, we selected six taxa with varying growth habits that spanned a wide phylogenetic and geographic breadth, had a documented history of invasion, and which had sufficient numbers of occurrence records in both native and introduced ranges to construct niche models (Table S4). These taxa are widely recognized as some of the most problematic non-native ferns: *Angiopteris evecta* (Forst.) Hoffm.*, Azolla filiculoides* Lam.*, Lygodium japonicum* (Thunb.) Sw.*, Macrothelypteris torresiana* (Gaudich.) Ching*, Salvinia molesta* D.Mitch., and *Sphaeropteris cooperi* (Hook. ex F.Muell.) R.M.Tryon. We downloaded occurrence data from GBIF using rgbif v3.8.1 (accessed July 2025; Chamberlain and Boettiger 2017; Chamberlain *et al*. 2025) and used CoordinateCleaner v3.0.1 (Zizka *et al*. 2019) to remove suspicious records. Additional erroneous records were inspected manually and removed as needed. For each taxon, we generated shapefiles for the native and introduced ranges from subsets of Natural Earth Data (https://www.naturalearthdata.com/) to crop a raster of all 19 bioclim variables available from WorldClim v2.1 (Fick and Hijmans 2017) at 30s resolution. To define the native range, we used the countries for which we verified the presence of the taxon. For the introduced range, we selected a focal introduced range and selected a conservative set of states/provinces/countries for which we verified the presence of the taxon (see Table S4 for list of native and invaded regions for each taxon). For taxa where there were two distinct invasions (i.e., *A. evecta, A. filiculoides, M. torresiana*), we also compared the climatic niches of the two invaded ranges.

We then used the scripts from Broennimann *et al*. (2012) to prepare the bioclimatic and occurrence data for analysis. The ‘PCAenv’ function was used in ecospat v4.1.2 (Di Cola *et al*. 2017) to generate a PCA of the climatic variables calibrated on both the native and introduced ranges. Broennimann *et al*. (2012) identified that ordination-based analyses out-performed species distribution models (SDMs), and that the PCAenv method is the most accurate method in niche overlap detection. Dynamics in the climatic niche were classified following the centroid, overlap, unfilling, and expansion (COUE) framework (Guisan *et al*. 2014). Briefly, changes in analog niche space (climate that is available in both the native and introduced ranges) can be described by measuring shift of the niche centroid, or based on their overlap (the intersection between native and invaded climatic niche space), stability (climatic niche space that remains constant or occupied in both ranges), expansion (climatic niche space that is occupied in the introduced range but not the native range-), and unfilling (climatic niche space that is occupied in the native range but not the introduced range) (Guisan *et al*. 2014). The overlap between the native and invaded climatic niches was calculated using Schoener’s D (Schoener 1970). We used a niche similarity test to compare the observed overlap metrics between the native and invaded niches to a distribution of overlaps generated between the observed invaded range and a randomized sampling of the native range and vice versa (Warren *et al*. 2008; Broennimann *et al*. 2012). We performed this tests with 1000 replicates, and set the following alternative hypotheses to test for niche conservatism between ranges: overlap.alternative = “lower”, expansion.alternative = “higher”, stability.alternative = “lower”, unfilling.alternative = “higher”. We further quantified dynamics in non-analog niche space (climatic nice space that is not available in both ranges) using scripts provided in Riera *et al*. (2025). Two additional metrics were calculated when examining niche dynamics in non-analog niche space: abandonment (climatic niche space that is occupied in the native range but not available in the introduced range) and pioneering (climatic niche space that is occupied in the introduced range but not available in the native range).

### Invasiveness, Polyploidy, and Genome Size

To test hypotheses about the interaction of genome size, ploidy, and invasiveness, we first downloaded the genome size and chromosome count database generated by Fujiwara *et al*. (2023). After filtering for taxa that had both genome size and ploidy data, there were very few non-native taxa, and therefore we searched the primary literature for data on genome size and chromosome counts for the non-natives in our checklist. For this extended dataset, the ploidy for each taxon was determined by comparing chromosome counts with the base chromosome count in the genus. Given that taxa often had multiple ploidal levels, we generated two datasets that retained either the lowest ploidy of each taxon (“low”) or the highest ploidy of each taxon (“high”). In these datasets, we only used a published genome size if the genome size estimate was directly estimated from an individual with that ploidy (Tables S2, S3). To place our results in a phylogenetic context, we again used the FTOL v1.7.0 (Nitta *et al*. 2022) and trimmed the phylogeny to only include the taxa that were present in our extended genome size and ploidy dataset. We then used a phylogenetically-informed generalized least square regression (pGLS) with a Brownian motion covariance matrix to assess the relationship between ln-transformed 2C genome size and invasive status using both the “low” and “high” datasets in the R package nlme v3.1-168 (Pinheiro *et al*. 2025). Tukey’s post-hoc tests were implemented using the ‘glht’ function in multcomp v1.4-28 (Hothorn *et al*. 2008). Unlike the continuous genome size, ploidy is a discrete variable which violates the distribution assumptions of a pGLS. Therefore, to assess the relationship of ploidy and invasive status, we used a phylogenetically-informed generalized linear mixed model (pGLM) for both “low” and “high” datasets with the R package phylolm v2.6.5 (Ho *et al*. 2024) using the “poisson_GEE” method. In order to assess pairwise comparisons among groups, we performed four tests with a different baseline for each test (i.e., native, casual, naturalized, invasive) and adjusted the raw *P-*values for multiple comparisons with a Bonferroni correction.

## RESULTS

### Dataset Compilation

Our initial list of non-native ferns (the union of three databases) included 269 candidate taxa (out of 11,916 species recognized in PPG I [2016], ca. 2.25%). The putative introduced ranges of many of these taxa could not be confirmed or had erroneous records, and were therefore removed. Our final checklist includes 83 taxa (ca. 0.69% of all fern species), including two subspecies of *Pteris cretica* L. Of these, we classified 18 as casual, 35 as naturalized (but not invasive), and 30 as invasive (Fig. 1, Table S1).

**Figure 1.**
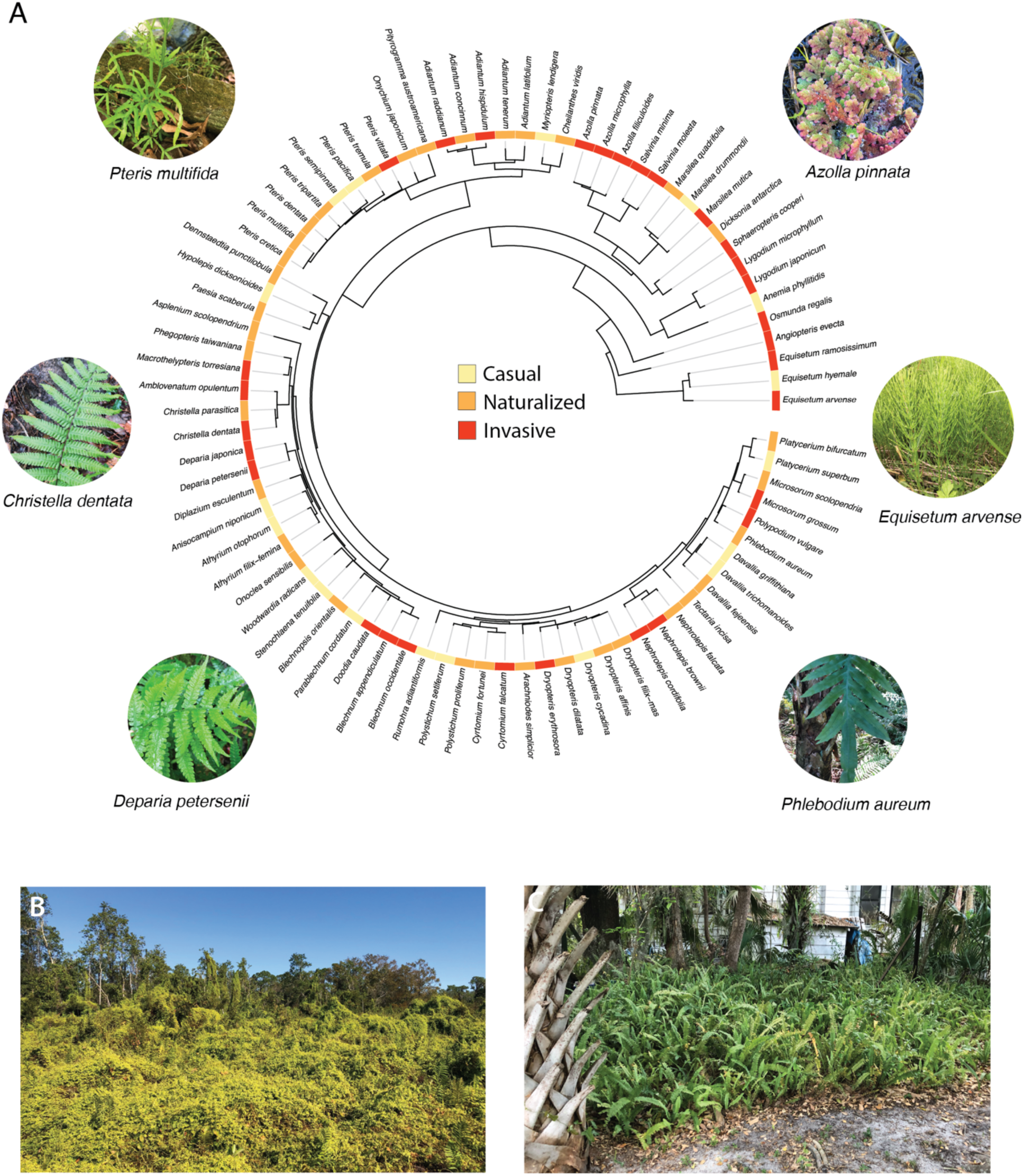
**A)** Phylogeny of non-native ferns subset from the Fern Tree of Life (Nitta et al., 2021). Species are classified as casual (yellow), naturalized but not invasive (orange), and invasive (red) based on the criteria listed in Table 1. Images of select non-native taxa from the root of the tree, counter-clockwise are *Equisetum arvense*, *Azolla pinnata, Pteris multifida, Christella dentata, Deparia petersenii,* and *Phlebodium aureum*. Examples of fern invasions of **B)** *Lygodium microphyllum* and **C)** *Neprolepis cordifolia* in Florida, USA. All images taken by J.A. Pelosi.

### Phylogenetic Distribution

A total of 21 families and 47 genera contained non-native fern taxa (Fig. 1, Tables S1, 5). Of these, 10 families and 16 genera had casual taxa, 14 families and 26 genera had naturalized taxa, and 14 families and 21 genera had invasive taxa. The best represented families are Pteridaceae, Dryopteridaceae, and Blechnaceae, while the genera with the greatest number of species are *Pteris* (Pteridaceae), *Adiantum* (Pteridaceae), and *Dryopteris* (Dryopteridaceae).

For the least restrictive (trait = all non-native categories) *D* statistic calculation (Fritz & Purvis 2010), we found no significant phylogenetic clustering and the distribution of the non-native trait was not significantly different from a random walk (*D*= 0.923, *P*Brownian = 0, *P*Random = 0.067, Table 2). For the moderately restrictive calculation (trait = naturalized + invasive status), we found statistically significant but weak phylogenetic clustering, with trait distribution significantly different from both a random distribution and Brownian motion (*D*= 0.862, *P*Brownian = 0, *P*Random = 0.01). For the most restrictive calculation (trait = invasive), we also found no significant phylogenetic clustering (*D*= 0.887, *P*Brownian = 0, *P*Random = 0.102). A total of 907 tips had significant positive LIPA values, indicating that these tips contributed to driving the phylogenetic signal we observed. Taxa with the highest LIPA values were clustered in Equisetaceae (three significant tips), Lygodiaceae (one), Salviniaceae (four), Marsileacaee (three), Pteridaceae (nine), Thelypteridaceae (three), Blechnaceae (five), Athyriaceae (three), Dryopteridaceae (four), and Nephrolepidaceae (two).

**Table 2.** Phylogenetic clustering (for binary traits) and phylogenetic signal (for continuous traits) for non-native ferns. *D* values near 0 represent a conserved trait, and values near 1 represent random evolution. Blomberg’s K or Pagel’s λ values near 1 indicate trait evolution consistent with Brownian motion. K values less than 1 or Pagel’s λ near zero indicates less phylogenetic signal than expected by Brownian motion. In all metrics, *P* values indicate statistical significance.

| *Phylogenetic Clustering* |  |  |  |  |  |
| --- | --- | --- | --- | --- | --- |
|  | Categorization Scheme | *D* Value | *P* (Brownian) | *P* (Random) |  |
|  | Naturalized & Invasive | 0.8616137 | 0 | 0.01 |  |
|  | Casual, Naturalized, & Invasive | 0.9232787 | 0 | 0.067 |  |
|  | Invasive | 0.8868383 | 0 | 0.102 |  |
| *Phylogenetic Signal* |  |  |  |  |  |
| | Categorization Scheme | Pagel's $\lambda$ | Blomberg's *K* | *P* (Pagel's $\lambda$) | *P* (Blomberg's *K*) |
|  | continuous - whole tree | 0.385474 | 0.0001506831 | 2.43143x10<sup>-42</sup> | 0.9190809 |
|  | continuous - only non-native species trimmed phylogeny | 0.231716 | 0.02480426 | 0.162444 | 0.05678864 |

Using the whole FTOL with both native and non-native taxa, we found significant phylogenetic signal for invasiveness with Pagel’s λ (Pagel 1999) (λ = 0.385, *P* = 2.43x10^-42^) but not Blomberg’s K (Blomberg *et al*. 2003) (K = 0.0001, *P* = 0.919). Blomberg’s K or Pagel’s λ values near 1 indicate trait evolution consistent with Brownian motion. In contrast, K less than 1 or Pagel’s λ near zero suggests less phylogenetic signal than expected by Brownian motion. Here, our λ value indicates a very weak but significant phylogenetic signal, while the K value is nonsignificant and suggests less signal than expected by Brownian motion.

With the trimmed FTOL with only non-native species, we found no significant signal for invasiveness with Pagel’s λ (Pagel 1999) (λ = 0.232, *P* = 0.162) and marginally significant signal with Blomberg’s K (Blomberg *et al*. 2003) (K = 0.025, *P* = 0.057). Our λ value here indicates no significant phylogenetic signal and that species are less similar in invasiveness than predicted by Brownian motion. Our Blomberg’s K value is marginally significant but again suggests less signal than expected by Brownian motion.

### Geographic Distribution

Regions differed significantly in the fractions of their fern flora that were sources (χ^2^ = 294.15, df = 8, *P* = 7.22 x10^-59^) and recipients (χ^2^ = 703.70, df = 8, *P* =1.14 x 10^-146^) of non-native ferns (Fig 2). A greater than expected proportion of the fern floras of Central/Temperate Asia, Australia, Europe, and the Pacific Islands are non-native in other regions (i.e., are over-represented as source regions, χ^2^ residual > 2). A lower than expected proportion of fern floras of Central America and South America are non-native in other regions (i.e., are under-represented as source regions, χ^2^ residual < -2). Geographic regions with a greater proportion of non-native ferns (i.e., recipient regions) than expected included North America, the Pacific Islands, Europe, and Australia. In contrast, Central America, South America, Central/Temperate Asia, and Eastern Asia had a lower than expected proportion of non-native ferns in their fern floras (i.e., under-represented recipient regions).

**Figure 2.**
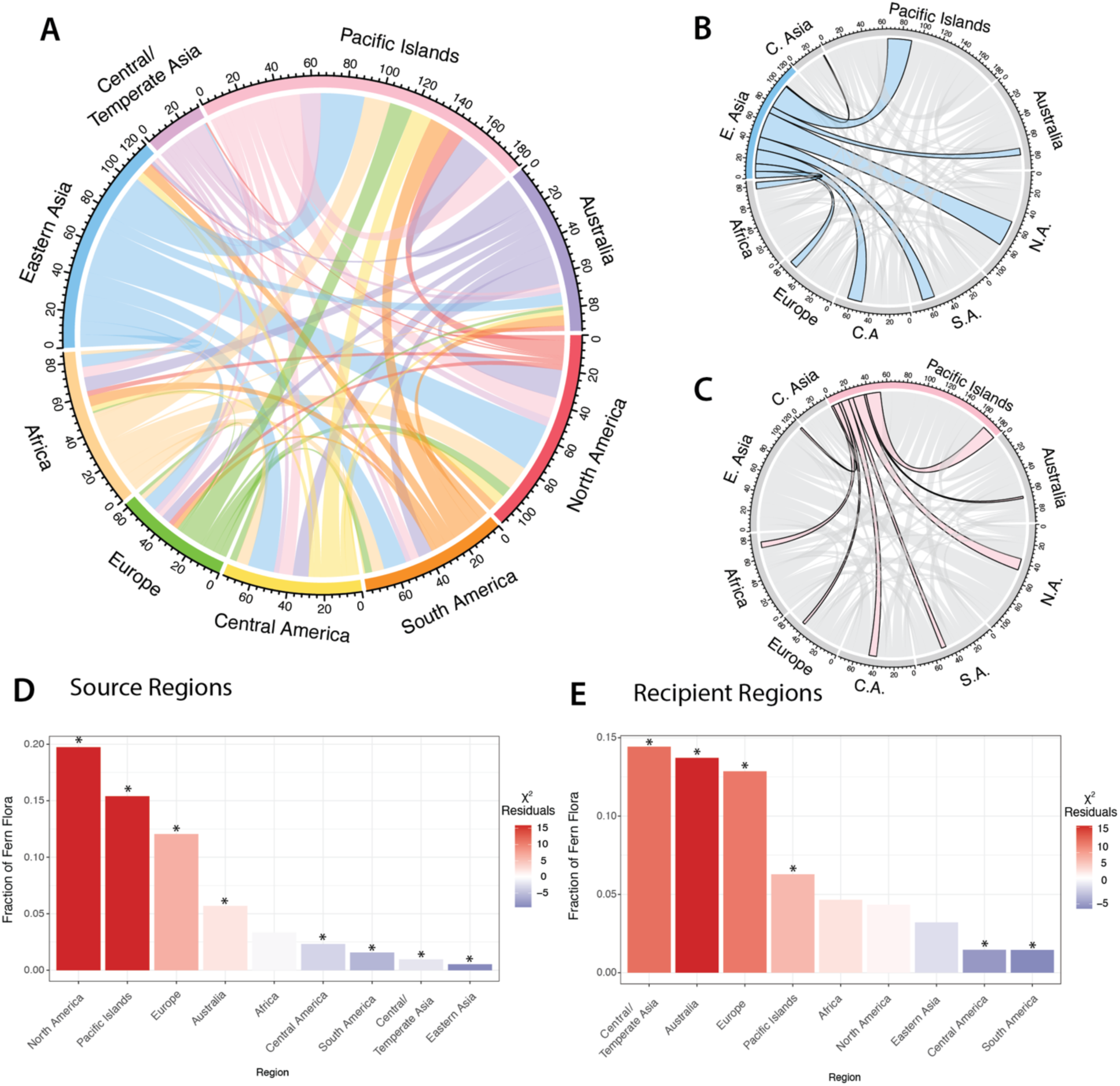
Chord diagrams showing the source and recipient geographic regions of non-native ferns, where chords connecting regions are proportional to the number of non-native taxa moving between those regions. **A)** Overall distribution of non-native fern taxa. **B)** and **C)** highlight particularly noteworthy regions. The fraction of fern floras in each geographic region that are non-native elsewhere (**D,** Source Regions) or are non-native to that region (**E,** Recipient Regions). Bars in **D** and **E** colored based on χ^2^ residuals, and regions with residuals with |χ^2^| > 2 are denoted with an asterisk. N.A. = North America, C.A. = Central America, S.A. = South America, E. Asia = Eastern Asia, C. Asia = Central/Temperate Asia.

### Climatic Niche Shifts

On average, Schoener’s *D* statistic, a measure of niche overlap that ranges from 0 (no overlap) to 1 (complete overlap) was low (average 0.1075; range = 0.0023-0.5334) when comparing the native and invaded climatic niches. In all cases, however, values of Schoener’s D (overlap) were not significantly lower than a distribution of overlaps generated between the observed invaded range and a randomized sampling of the native range (*P* > 0.05, Table S4). While Schoener’s D is an overall look at niche overlap, differences between the climatic niches can be broken down into specific dynamics (Fig. 4). When comparing only analog niche space, we measured dynamics of niche expansion, stability, and unfilling. The amount of expansion in the invaded range compared to the native range was small (weighted proportion of climatic niche space in this category: mean = 0.0736; range = 0-0.2860) and was not significant for any comparison (*P >* 0.05). In contrast, the proportion of the niche that was stable was high (mean = 0.9263; range = 0.7140-1) and was not significantly less than expected (*P >* 0.05). Niche unfilling was moderate across comparisons and highly variable (average 0.4806; range = 0.0803-0.8791), although these were also all non-significant (*P* > 0.05). When comparing both analog and non-analog niche dynamics, we measured metrics of abandonment and pioneering in addition to expansion, stability, and unfilling, relative to the entire niche space available. Niche abandonment was relatively common, with a mean of 0.2243 (range= 0-0.3941), while pioneering was only observed in two taxa (0.5731 for *Salvinia molesta* and 0.785 for *Sphaeropteris cooperi*).

**Figure 3.**
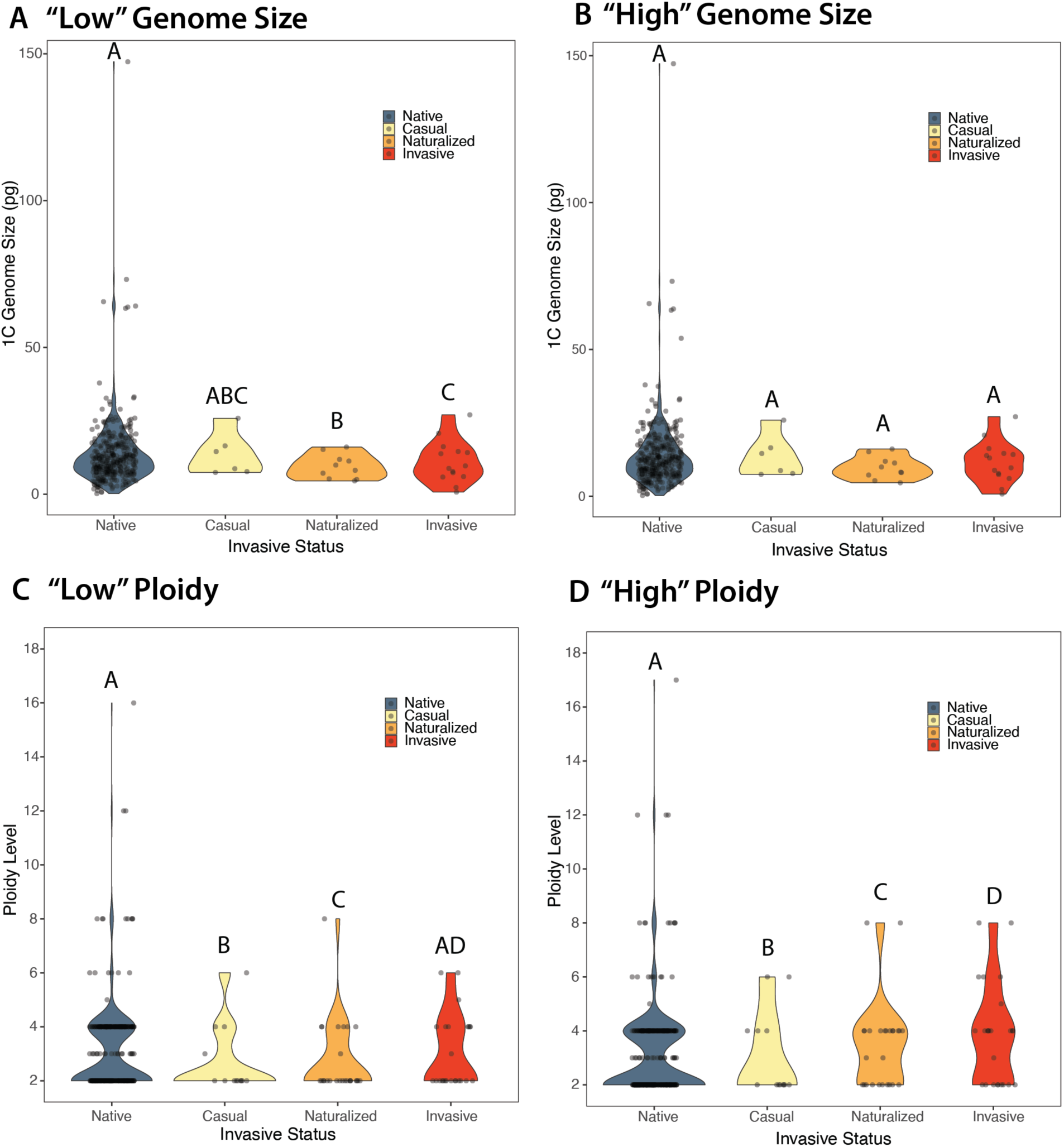
Association between genome size (**A, B**) and ploidy level (**C, D**) for “low” (**A, C**) and “high” (**B, D**) datasets, which retained the lowest or highest value, respectively, for species with variable ploidies. Results of pGLS ANOVAs are displayed for each dataset, and significance for pairwise post-hoc tests are shown with letters.

**Figure 4.**
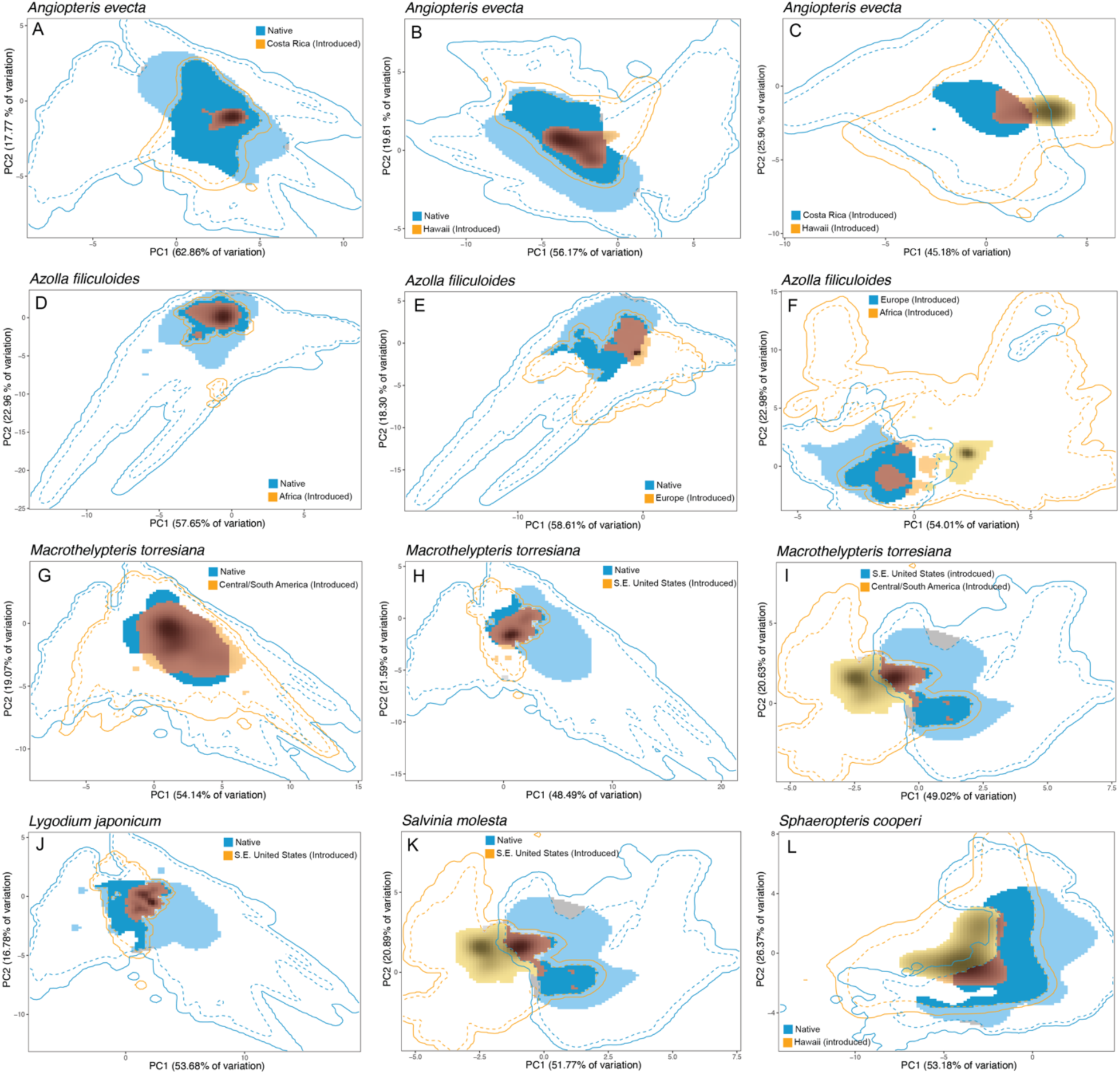
Climatic niche dynamics summarized on the first two principal component axes. In each graph, the solid blue line denotes the total climatic space available in the native range while the solid orange line denotes the total climatic space available in the invaded range. Dashed lines represent 75% of the available climate space. In analog niche space, solid red coloration shows stability (climatic niche space that is occupied in both the native and invaded ranges), orange shows expansion (climatic niche space that is occupied only in the introduced range), and blue shows unfilling (climatic niche space that is occupied only in the native range). In non-analog niche space, light blue coloration shows abandonment (climatic niche space that is occupied in only the native range and not available in the introduced range), and yellow shows pioneering (climatic niche space that is occupied only in the invaded range and not available in the introduced range). The density of occurrences depicted by the brightness (e.g., darker indicates higher occurrence density). **C, F,** and **I** are comparisons of distinct invasions for *Angiopteris evecta, Azolla filiculoides,* and *Macrothelypteris torresiana*, respectively.

When comparing different invaded ranges of *Angiopteris evecta, Macrothelypteris torresiana*, and *Azolla filiculoides,* the average Schoener’s D metric was similar to the average for the native to invaded range comparisons (mean = 0.1405; range = 0.0020-0.3464). These overlap values were not significantly lower than a random distribution (*P* > 0.05). The niche expansion factors for these comparisons were much greater than the native-invaded comparisons (mean = 0.4061; range = 0.3458-0.4771), while the stability factor was lower (mean = 0.5939; range = 0.5229-0.6542). The unfilling factor was slightly greater than the native-invaded range comparisons (mean = 0.62; range = 0.4359-0.7292). The niche expansion, stability, and unfilling factors for the invaded-invaded comparisons were also non-significant (*P* > 0.05). In non-analog niche space, abandonment from the first invaded niche was relatively low (mean = 0.0362; range = 0.0007-0.1090), while pioneering in the second niche was variable and high (mean = 0.5522; range = 0.1489-0.9349).

### Association of Genome Size, Ploidy, and Invasiveness

We collected ploidy data for 420 fern species (356 native, 13 casual, 26 naturalized, 25 invasive), and genome size data for 385 fern species (356 native, 6 casual, 10 naturalized, 13 invasive) (Tables S2, S3). For the “low” 2C genome size dataset (where we retained the lowest genome size reported for taxa with multiple ploidy levels), there was a significant relationship between invasive status and ln(2C genome size) (pGLS ANOVA *P* = 0.001). Within this test, the 2C genome sizes of naturalized taxa (mean = 18.97 pg) and invasive taxa (mean = 23.00 pg) differed significantly (*P =* 0.001), as did those of naturalized taxa and native taxa (mean = 26.89 pg; *P* = 0.026) and invasive and naturalized taxa (*P =* 0.001). No other pairwise differences were significant (all *P* > 0.05; Fig. 3, Table S6). For the “high” 2C genome size dataset (where we retained the highest genome size reported for taxa with multiple ploidy levels), we found no significant relationship between 2C genome size and invasive status (pGLS ANOVA *P* > 0.05; Fig. 3, Table S6). As such, we found no significant differences between any of the invasive status pairings (native mean = 27.29 pg, casual mean = 26.96 pg, naturalized mean = 19.54 pg, invasive mean = 23.00 pg; all *P* > 0.05). For the ploidy datasets, comparisons among the different categories were consistently different, although the direction of the difference varied between the “low” and “high” datasets. From the pGLM of the “low” ploidal level dataset, the only non-significant difference in ploidal level was between native (mean = 3.01) and invasive taxa (mean = 3.04; Bonferroni-corrected *P* = 0.476). We did find significant differences when comparing invasive (mean = 3.04), casual (mean = 2.69), and naturalized (mean = 2.73) taxa (all Bonferroni-corrected *P* < 0.05). For the high ploidy dataset, our results were nearly the same, although the direction of the differences was flipped (taxa further along the invasion continuum had higher ploidal levels) and all comparisons were significant (all Bonferroni-corrected *P <* 0.05). In this dataset, the mean ploidal levels were 3.09 for native taxa, 3.08 for casual taxa, 3.46 for naturalized taxa, and 3.84 for invasive taxa.

## DISCUSSION

Non-native ferns represent a non-trivial component of the world’s exotic floras and can have significant ecological and economic impacts in their introduced ranges (Robinson *et al*. 2010; Jones *et al*. 2019). Despite their importance, their study remains underrepresented compared to other plant groups (Pyšek *et al*. 2017), and available information on non-native species and their progression along the invasion continuum is often fragmented and restricted to particular taxa (Pemberton *et al*. 2002; Lott *et al*. 2003; Murakami *et al*. 2007; Chau *et al*. 2013; Yañez *et al*. 2023) or regions (Wilson 2002; Yañez *et al*. 2020). The results presented here provide a comprehensive global review and classification of non-native ferns and offer a framework to test hypotheses about their colonization processes.

### Overview of dataset

Based on an extensive literature review, we identified 83 non-native fern species worldwide from an initial pool of 269 potential taxa compiled from major databases—including World Ferns (Hassler 1994-2026, accessed 11 May 2021), the Global Naturalized Alien Flora (GloNAF; van Kleunen et al. 2019), and the list of invasive terrestrial ferns assembled by Jones *et al*. (2019). Because Jones *et al*. (2019) was largely based on datasets from North America, Europe, Asia, and Australia, but did not include information from South America, we placed special emphasis on gathering regional studies documenting non-native ferns in this region. A major challenge during data compilation was confirming native versus introduced ranges, as information on species origins and human-mediated dispersal pathways is frequently incomplete or dispersed across local floras, systematic revisions, and ecological studies. The considerable difference between our initial inventory and the final dataset underscores the importance of contrasting global databases with detailed regional and local sources that provide verified specimen records and biogeographic context.

In addition, the invasion-stage classification assigned here differs slightly from that proposed by those authors for 32 of the species on our list, despite the use of comparable criteria (Pyšek et al. 2004; Falk-Petersen et al. 2006; Blackburn et al. 2011). For 13 species, our review revealed more advanced stages in the invasion continuum, aided by detailed information (often contained in descriptive notes or discussions) found in taxonomic treatments, ecological analyses, and regional floras, which provide insights into species’ distributions in their introduced ranges. Several species identified as invasive in our analysis had already been noted by Robinson (2010) and Akomolafe and Rahmad (2018) as problematic due to their ability to spread, displace native species, or negatively affect natural habitats and human activities. While these studies primarily focused on the environmental problems caused by these species, they did not interpret their establishment and spread within an invasion ecology framework—an aspect to which this article contributes novel synthesis of information.

### Result I. Weak or no phylogenetic clustering of non-native ferns

Invasiveness may not be randomly distributed across the plant tree of life. Traits associated with invasion success, such as reproductive strategies, dispersal mechanisms, physiological tolerances, or competitive ability, could be phylogenetically conserved, meaning they are more likely to be shared by closely related species (Ackerly 2009). If there is phylogenetic clustering of invaders, this could help identify conserved traits associated with invasiveness and be used to predict which other species may become invasive and where to focus mitigation efforts (Pyšek 1998; Lambdon 2007). Studies on angiosperm invaders have found significant, but often weak or moderate, phylogenetic clustering (Pyšek 1998; Cadotte *et al*. 2009; Pertierra *et al*. 2023). Pyšek *et al*. (2015) identified specific conserved biological traits (including seed bank persistence, longer flowering periods, clonality, and plant height) associated with naturalization success. Additionally, Cadotte *et al*. (2009) found that successful large-scale invaders tended to be phylogenetically related to other successful invaders with which they share ecological traits that promote invasion. Notably, Qian *et al*. (2022) found that phylogenetic clustering was greater in more harmful invaders than in introduced species as a group

In the ferns, Jones *et al*. (2019) found that five families (Psilotaceae, Nephrolepidaceae, Equisetaceae, Onocleaceae and Lygodiaceae) were over-represented in their dataset of non-native terrestrial ferns. Using a phylogenetic framework, our analyses provide a nuanced perspective that largely agrees with the findings in Jones *et al*. (2019). We recovered no or very weak phylogenetic clustering of invasive status in ferns. When considering the entire phylogeny (i.e., all native and non-native taxa), we found weak but significant phylogenetic signal for invasive status as a continuous trait. Similarly, when we considered a binary trait of naturalized or invasive status versus all other fern taxa, we recovered very weak, but significant clustering. In particular, the families driving this phylogenetic signal largely overlapped with Jones *et al*. (2019), including Equisetaceae, Lygodiaceae, Nephrolepidaceae, and the two clades of heterosporous water ferns (not included in Jones *et al*. 2019), Salviniaceae and Marsileaceae. Overall, our analyses found that phylogenetic clustering was only significantly different from random when considering naturalized and invasive species together, but not invaders alone, unlike the stronger relationships of invaders in particular found by Qian *et al*. (2022).

Given our results, invasiveness in ferns appears to be dictated by a mixture of conserved and non-conserved traits, and which traits dictate invasion success may differ across clades, regions, and ecosystems. For example, traits that facilitate invasion in floating water ferns are likely to differ from those in terrestrial invaders. Examining phylogenetic clustering at smaller phylogenetic scales may be more informative. As evidence of this, Pertierra *et al*. (2023) found that the phylogenetic distribution of invasiveness varies among major subclades of Poaceae. Additionally, their study highlights a positive relationship between invasiveness and diversification rates, suggesting that greater evolutionary versatility and radiation within a particular clade could promote the emergence of new invasive species. A clear understanding of the phylogenetic relationships among non-native fern taxa would further aid in the prediction and mitigation of future invasions.

### Result II. Some regions are overrepresented as sources and recipients of introductions

Understanding where non-native plants originate and where they accumulate is a fundamental task in invasion biology. While the “imperialist dogma” suggests that European species should be over-represented in recipient regions due to extensive and long-lasting colonization (Crosby and Worster 2004), mounting evidence indicates that there is substantial temporal variation in source-recipient dynamics, with additional regions increasingly recognized as sources (Seebens *et al*. 2018). The complexities of human movement are important factors to consider in analyzing such data (Hulme 2009) and global trade routes are known to be correlated with non-native species establishment (Chapman *et al*. 2017). Furthermore, the effort expended on surveillance of non-natives varies greatly between countries. For example, New Zealand and Hawaii employ comprehensive programs to detect and monitor non-native species, potentially resulting in an overrepresentation of such locations as recipient regions (e.g., McDonald *et al*. 2020; Bird *et al*. 2025).

Keeping those important sources of variation in mind, we identified several global regions that were over-represented as sources of non-native ferns identified to date: Central/Temperate Asia, Australia, Europe, and the Pacific Islands. Among recipient regions, North America, the Pacific Islands, Europe, and Australia were over-represented. These patterns differ partially from global patterns of non-native distributions for all plants, in which North America has been the overall largest recipient, and continents in the Northern Hemisphere have been the largest donors (van Kleunen *et al*. 2015). This may be due in part to a particular historical fascination with ferns during the Victorian era in the horticultural trade of Europe and its colonies termed “pteridomania” (Allen 1969; Boyd 1992, 2005; Whittingham 2009). Nurseries in the U.K. imported fern taxa from around the world, with which collectors traveled, showcasing wild and horticultural hybrids globally (Boyd 2005). This fascination with ferns in the mid-nineteenth century likely facilitated the spread of non-native ferns globally, especially among areas colonized by the British Empire. For example, Australia, Europe, and the Pacific Islands (all regions colonized by Britain) were both over-represented sources and recipients of non-native fern taxa, and collection and transportation of ferns from Asia, an over-represented source of non-native ferns, was facilitated by the East India Company (Keogh 2023). Although the fern craze of Victorian Britain ended in the early twentieth century, McCulloch-Jones *et al*. (2021) found that current sale and trade of ferns through physical and e-commerce nurseries is associated with the probability of establishment.

Importantly, there were several taxa for which it was difficult for us to fully ascertain their “native” geographic range (e.g., *Pteris vittata* L.*, Macrothelypteris torresiana*). Because these taxa are cosmopolitan, they tend to be collected less frequently than others, and their representation in herbaria is therefore underestimated relative to their true geographic distribution. Also, when their broad distribution is partly driven from cultivation as ornamentals in garden settings (Robinson *et al*. 2010; Pemberton 2025), the boundaries between native and introduced ranges become blurred and tracing the origins of such plants can be especially challenging without in-depth studies. Furthermore, signatures of admixture and multiple introductions (possibly from different sources), which are common in many invasions (Dlugosch and Parker 2008), may obscure clear geographic patterns but represent true biological complexities. In many cases determining the source population/region for taxa may require extensive population-level sampling and population genetic analyses.

### Result III. Climatic niches are often conserved between native and introduced ranges, but can differ between introductions

Comparing the niches occupied between native and invaded ranges of a species can reveal whether their native niche predicts further expansion into new areas of an introduced range, and whether invaders are problematic in part because they are able to expand their niche beyond predictions from the native range (Petitpierre *et al*. 2012). Therefore, the ability to predict the distribution of non-native species from their climatic niche relies on climate matching between ranges (Richardson and Thuiller 2007; Thuiller *et al*. 2008). If species explore novel niche space in the non-native range, niche models will be unable to accurately forecast their spread, and this problem appears to be common among invasive plants (Atwater *et al*. 2018). Several studies of invasive plants have found that niche shifts are common (Gallagher *et al*. 2010; Early and Sax 2014; Atwater *et al*. 2018), while others had contradictory findings that shifts in climatic niches were rare (Petitpierre *et al*. 2012; Liu *et al*. 2020), with others such as Riera *et al*. (2025) suggesting that conservation of the niche is largely dependent on native niche breadth. The most comprehensive study by Atwater *et al*. (2018), which included seven fern taxa, found evidence of climatic niche shifts in 65–100% of 815 invasive terrestrial plant species in different regions, and that many of those shifts were associated with non-analog niche space: abandonment of climatic space available in the native but not in the invaded range accounted for 23% of niche dynamics, while pioneering (occupying niche space only available in the invaded range) accounted for 10% of niche dynamics.

The niches of native and invaded ranges have not been compared for ferns. Some authors have begun characterizing the niches of invasive ferns to predict their spread. For example, Goolsby (2004) and Christenhusz and Toivonen (2008) used ecological niche modeling to predict the potential spread of *Lygodium microphyllum* and *Angiopteris evecta,* respectively. A broader study of six horticulturally-traded ferns used niche modeling to generate risk assessments based on the assumption of niche stability (McCulloch-Jones *et al*. 2023). All of these studies predicted further spatial expansion under the assumption of niche stability and climate matching between ranges (i.e., no change in the niche occupied), but this assumption and the possibility of ferns exploring novel niche space has not been tested.

For four of the six fern species that we examined, the climatic niches in their introduced range largely overlapped with those of their native range. The climatic niches for the introduced ranges of *Angiopteris evecta, Azolla filiculoides, Macrothelypteris torresiana,* and *Lygodium japonicum* were nearly entirely within their native range climatic niche space with very little evidence of expansion into analog niche space or pioneering in non-analog niche space. In contrast, *Salvinia molesta* and *Sphaeropteris cooperi* both pioneered non-analog niche space while unfilling analog niche space. Pioneering in *Salvinia molesta* is particularly prominent; over half of the niche dynamics were classified as the exploration of non-analog niche space. Although we analyzed only a subset of invasive taxa, there does not appear to be a clear relationship between niche dynamics and growth habit or phylogenetic position in these species. For example, both *A. filiculoides* and *S. molesta* are floating water ferns, yet exhibit profoundly different patterns: the climatic niche dynamics of *A. filiculoides* are characterized by high stability, while those of *S. molesta* are largely driven by pioneering. Notably, two of the species that we examined (*Lygodium japonicum* and *Sphaeropteris cooperi*) were included in the risk assessments of McCulloch-Jones *et al*. (2023); while these analyses included occurrences from both native and invaded ranges, our finding that there is pioneering in the invaded range of *S. cooperi* indicates that the extent of further niche expansion is possible, which would put additional areas at risk.

While climatic niche space is largely conserved between native and introduced ranges of these ferns, we found that comparing distinct invasions of the same species paints a more dynamic picture of climatic niche patterns. For example, when comparing the climatic niche space of *Macrothelypteris torresiana* in its introduced range in the southeastern United States to that in Central and South America, there was relatively little niche overlap and there was exceptional pioneering in the latter range. Differences in the dynamics between invasions have been reported in other systems (e.g., *Centaurea stoebe,* Broennimann *et al*. 2014) and may be associated with differences in the invasion process such as residence time, introduction pathway, site of origin, and genotypes introduced (Richardson and Pyšek 2012; Gioria *et al*. 2023). Notably, different biotic interactions can also alter climatic niches in different regions, and while ferns are typically thought to largely lack herbivores, recent work has contradicted this assumption and highlighted a wide variety of fern-insect interactions (Hendrix and Marquis 1983; Mehltreter 2010; Suissa *et al*. 2024; Cariglino *et al*. 2025). Contrasting niche occupancy patterns across native and multiple invaded regions offers further opportunities to identify how biotic and abiotic variables shape fern distributions.

### Result IV. Naturalized ferns have smaller genomes

Numerous studies of angiosperms have demonstrated that those considered weedy, whether locally or regionally, have significantly lower amounts of nuclear DNA and smaller genome sizes than other plants (Bennett *et al*. 1998; Pandit *et al*. 2014; Suda *et al*. 2015; Pyšek *et al*. 2023). Smaller genome sizes are associated with life cycle characteristics common in colonizing plants such as (1) establishing quickly and displaying a short juvenile phase; (2) developing rapidly throughout the life cycle and having a short minimum generation time; and (3) rapidly producing many small propagules (Oka and Morishima 1982; Rejmánek and Richardson 1996). The timing of developmental stages in the life cycle is closely tied to the timing of the cell cycle. Lower DNA content has been linked to shorter cell cycles and higher cell division rates (Gregory 2001). The large genome constraint hypothesis (Knight *et al*. 2005) posits that such functional traits (e.g., propagule mass and number, specific leaf area, relative growth rate, and generation time, Suda *et al*. 2015) are associated with genome size and that large genomes are costly since they directly negatively impact these traits. Given that many of these traits are associated with invasion success (Suda *et al*. 2015), it follows that genome size should therefore have a negative relationship with invasion success, and analyses of angiosperm datasets support this hypothesis that many invaders have smaller genomes than their native counterparts (Chen *et al*. 2010; Kuester *et al*. 2014; Pandit *et al*. 2014; Suda *et al*. 2015). For example, Suda *et al*. (2015) recovered that the median genome size of invasive angiosperms was nearly half that of native species and Pandit *et al*. (2014) found a strong significant negative relationship between haploid genome size and invasiveness. To date, ferns have been excluded in these studies in part due to the lack of a comprehensive checklist of non-native ferns, therefore it is unclear if invasive status is correlated with genome size in this lineage.

In our dataset, we also found that genome sizes tended to be smaller in non-native ferns. Although significant, this difference was not as substantial as observed in angiosperms. For our “high” genome size dataset (retaining the highest estimates when reports of genome size were variable) the average native fern 2C genome size was 26.89 pg (mean 2C approximately 26.5 Gb, or 1C approximately 13.2 Gb), which is similar to previous estimates over all ferns (Nakazato *et al*. 2008; Sessa and Der 2016). Naturalized, but not invasive, taxa had the smallest genomes (mean 2C = 18.97 pg), while invasive ferns had significantly smaller genomes than native ferns (mean 2C = 23.00 pg). In contrast to the >50% smaller genome size of invasive angiosperms reported by Suda *et al*. (2015, average native 1C = 5.83 pg, average invasive 1C = 2.39 pg), we found that naturalized ferns had genomes that were about 70% the size of native fern genomes, and invasive species had genomes that were approximately 85% the size of native fern genomes. Such a difference in the relationship between genomic content and non-native status could highlight fundamental differences in the underlying genomic biology of ferns compared to seed plants. For example, fern lineages tend to have larger genome sizes and less genome reduction after whole genome duplication than angiosperms over evolutionary time (Haufler and Soltis 1986; Haufler 1987; Barker 2009) such that there may be less opportunity for smaller genomes to predominate among successfully introduced ferns.

Our results are also consistent with a previous analysis that suggested that the relationship of genome size and invasion stage was not linear. In their analysis of naturalized seed plants, Pyšek *et al*. (2023) found a non-linear (quadratic) relationship between genome size and naturalization and invasion extent, such that small genomes were associated with naturalization success (intermediate state) but not native or invasive status. Interestingly, this relationship suggests that small genomes play a role in facilitating the naturalization process, but might constrain their spread (Pyšek *et al*. 2023). We also found that the genome size of naturalized, but not invasive, plants were smaller than those that were classified as invasive. The results suggest that trait values influenced by genome size are not uniform across the invasion continuum. Notably, genome sizes can also evolve within a species during the process of invasion, complicating these types of analyses (Lavergne *et al*. 2010; Cang *et al*. 2024). Our dataset does not have the resolution to compare genome sizes among populations within a species, and investigations of genome size evolution in the invaded ranges of non-native ferns are lacking. Such studies present a valuable area for further research in ferns.

### Result V. Invaders have higher ploidal levels

Polyploidy, the multiplication of the entire set of genomic material within an organism, is a widespread phenomenon among plants (Ledyard Stebbins 1940; One Thousand Plant Transcriptomes Initiative 2019) with profound ecological and evolutionary effects (Van de Peer *et al*. 2009, 2021). At the genome-level, additional gene copies provide material upon which selection can act (e.g. neo- and sub-functionalization of duplicate gene copies), alter gene regulatory networks, and can mask deleterious alleles (Li *et al*. 2021). Furthermore, polyploids (particularly allopolyploids which are a product of hybridization) may have fixed heterozygosity (Roose and Gottlieb 1976) and heterosis (East 1936), giving them a distinct advantage over diploids (Soltis and Soltis 2000). These genomic changes can be associated with ecological, physiological, and reproductive parameters that may drive the success of polyploids (reviewed by te Beest *et al*. 2012). For example, increased gene diversity through heterozygosity could allow increased phenotypic plasticity and therefore the ability to persist in the face of environmental changes (Leitch and Leitch 2008; te Beest *et al*. 2012). Given the numerous associations between polyploidy and functional traits, polyploidy has long been considered an important factor in invader success (e.g. Mulligan 1960, 1965; Baker 1965; Ehrendorfer 1965). Indeed, recent large-scale investigations of polyploidy and invasion have supported this hypothesis (e.g. Pandit *et al*. 2006, 2011; Moura *et al*. 2021; Pyšek *et al*. 2023).

The possibility of a relationship between polyploidy and invasion success in ferns, however, has yet to be tested. Nearly one-third of speciation events in ferns is associated with a change in ploidal level (Wood *et al*. 2009) compared to around 15% in angiosperms, and nearly all ferns have at least one whole genome duplication in their evolutionary history (Pelosi *et al*. 2022). Recent work studying the implications of polyploidy in ferns has uncovered that some allopolyploids experience niche expansion and occupy novel niche space compared to their diploid progenitors (Marchant *et al*. 2016; Wefferling *et al*. 2024), and that the niches of polyploid gametophytes tend to be more flexible than their haploid counterparts (Blake-Mahmud *et al*. 2025).

Our results interestingly revealed two contrasting patterns when using datasets that only took the lowest ploidal level (”low”) and the highest ploidal level (”high”) for each taxon when there were multiple estimates for a taxon. In the “low” dataset, we found that both casual (mean = 2.69) and naturalized (mean = 2.73) ferns have lower ploidal levels compared to native taxa (mean = 3.01), while there was not a significant difference between native and invasive (mean = 3.04) taxa. In contrast, the same analysis with the “high” dataset found that naturalized (mean = 3.46), and invasive (mean = 3.84) taxa had greater ploidy levels compared to native (mean = 3.09) taxa, while casual taxa had slightly lower ploidy levels (mean = 3.07). This latter result is largely congruent with previous findings in other plant lineages, where invasive plants tend to have higher ploidy levels (Pandit *et al*. 2006, 2011; Moura *et al*. 2021; Pyšek *et al*. 2023).

Notably, our ploidy data do not distinguish whether the ploidal level reflects native or invaded range populations or both, which may partly explain the contrasting trends between the “low” and “high” datasets. Few studies have explicitly examined the possibility of ploidy variation between native and introduced ranges or within the introduced range in ferns. For some taxa, only a single cytotype is represented in the invaded range, whereas multiple cytotypes may exist in the native range (e.g., *Lygodium japonicum*, Pelosi *et al*. 2023; *Phegopteris taiwaniana* T.Fujiw., Ogiso & Seriz., Fujiwara et al. 2021). In contrast, novel cytotypes may originate in the introduced range, although no evidence exists for this in the taxa we investigated. Across angiosperms, species that contain multiple cytotypes (e.g., diploid and polyploid) are more likely to be non-native than those that are only diploid (Pyšek *et al*. 2023). A detailed comparison of cytotype variation across the ranges of non-native ferns would provide additional resolution regarding the contributions of ploidy to the stages of fern invasions.

## Conclusions

This study integrated regional floras and natural history collections to compile a well-supported dataset of non-native ferns, which enabled tests of ecological, genomic, and phylogenetic features associated with the colonization of new habitats by ferns. In summary:

- We identified 83 strongly supported non-native fern species worldwide. Reliable evaluations of the distribution and invasion stage of non-native ferns depended on the systematic incorporation of regional and local sources—floras, taxonomic treatments, and ecological studies—which provided the verified records and contextual detail often missing from global databases.
- There was a weak but significant phylogenetic signal for invasive status across the fern phylogeny when treated as a continuous character or when including both naturalized and invasive status together as a binary “invader” trait. This differed somewhat from analyses of angiosperms that identified stronger phylogenetic clustering particularly for non-native plants with invader status.
- Central/Temperate Asia, Australia, Europe, and the Pacific Islands have acted as source regions of non-native species, contributing a greater-than-expected share of fern taxa to other regions, and, together with North America, also harbor higher-than-expected proportions of non-native ferns themselves.
- Introduced ferns generally occupied climatic niches that strongly overlapped with those of their native ranges, except in some cases (e.g., *Salvinia molesta, Sphaeropteris cooperi*) that showed notable pioneering into non-analog climatic space. Comparisons among independent invasions of the same species revealed marked differences in niche dynamics among invasions, underscoring that invasion history and local context shape climatic niche outcomes.
- Naturalized, but not invasive, fern taxa had the smallest genomes, while invasive ferns had significantly smaller genomes than native ferns. These differences were not as substantial as observed in angiosperms.
- Analyses using minimum vs. maximum ploidy values yielded contrasting patterns, but the “high” dataset of maximum values indicated that naturalized and invasive ferns tend to have higher ploidal levels than natives. This aligns with evidence across plants suggesting that polyploidy can facilitate invasion, though the specific mechanisms remain unclear.

## SUPPLEMENTARY INFORMATION

**Table S1.** List of non-native fern taxa with invasion continuum categorization, including references of regional and local floras and/or peer-reviewed publications and quotes from the primary literature, floras, or herbarium labels to inform and justify categorization.

**Table S2.** Table with published genome size if the genome size estimate was directly associated with a ploidal level for “high” genome size / ploidal level dataset.

**Table S3.** Table with published genome size if the genome size estimate was directly associated with a ploidal level for “low” genome size / ploidal level dataset.

**Table S4.** Results of ecological niche shift dynamics analyses.

**Table S5.** Comparisons between species studied in this article, (Robinson *et al*. 2010), (Akomolafe G. F. 2018), and (Jones *et al*. 2019).

**Table S6.** Ploidy and genome size comparison results.

## Author Contributions (CRediT)

**Jessie A. Pelosi:** Conceptualization; Data Curation; Formal Analysis; Supervision; Visualization; Writing - Original Draft Preparation; Writing - Review and Editing

**Agustina Yañez:** Conceptualization; Data Curation; Supervision; Writing - Original Draft Preparation; Writing - Review and Editing

**Leah N. Veldhuisen:** Formal Analysis; Visualization; Writing - Original Draft Preparation; Writing - Review and Editing

**Anthony J. Dant:** Formal Analysis; Visualization; Writing - Original Draft Preparation; Writing - Review and Editing

**Poppy C. Northing:** Formal Analysis; Visualization; Writing - Original Draft Preparation; Writing - Review and Editing

**Rebecca G.W. Bland:** Formal Analysis; Visualization; Writing - Original Draft Preparation; Writing - Review and Editing

**Weston L. Testo:** Conceptualization; Data Curation; Supervision; Writing - Original Draft Preparation; Writing - Review and Editing

**Katrina M. Dlugosch:** Conceptualization; Supervision; Writing - Original Draft Preparation; Writing - Review and Editing

## AVAILABILITY OF DATA AND MATERIALS

Code used for this study is available at: https://github.com/jessiepelosi/NaturalizedFerns.

## ACKNOWLEDGEMENTS

We thank Emily Sessa for advice on African floras and Melissa Smith for constructive conversations on invasive ferns.

